# LAMP3 is critical for surfactant homeostasis in mice

**DOI:** 10.1101/2021.02.05.429758

**Authors:** Lars P. Lunding, Daniel Krause, Guido Stichtenoth, Cordula Stamme, Niklas Lauterbach, Jan Hegermann, Matthias Ochs, Björn Schuster, Radislav Sedlacek, Paul Saftig, Dominik Schwudke, Michael Wegmann, Markus Damme

**Affiliations:** Airway Research Center North, German Center for Lung Research (DZL), Borstel, Germany; Division of Asthma Exacerbation & Regulation, Research Center Borstel, Leibniz Lung Center, Borstel, Germany; Bioanalytical Chemistry, Priority Research Area Infections, Research Center Borstel, Leibniz Lung Center, Borstel, Germany; Department of Pediatrics, University of Lübeck, Lübeck, Germany; Division of Cellular Pneumology, Research Center Borstel, Leibniz Lung Center, Borstel, Germany, and Department of Anesthesiology and Intensive Care, University of Lübeck, Lübeck, Germany; Institute of Biochemistry, Christian-Albrechts-University Kiel, Kiel, Germany; Institute of Functional and Applied Anatomy, Research Core Unit Electron Microscopy, Hannover Medical School, Hannover, Germany; Institute of Functional Anatomy, Charité Medical University of Berlin, Berlin, Germany; German Center for Lung Research (DZL), Berlin, Germany; Czech Centre for Phenogenomics, Institute of Molecular Genetics of the Czech Academy of Sciences, Vestec, Czech Republic; German Center for Infection Research (DZIF), TTU Tuberculosis, Borstel, Germany

**Author notes:** Corresponding authors: Markus Damme, Christian-Albrechts-University Kiel, Institute of Biochemistry, Olshausenstr. 40, 24098 Kiel Germany; Mail, Michael Wegmann, Division of Asthma Exacerbation & Regulation, Priority Area Asthma & Allergy, Research Center Borstel- Leibniz Lung Center, Borstel, Germany. Mail.

## Abstract

Lysosome-associated membrane glycoprotein 3 (LAMP3) is a type I transmembrane protein of the LAMP protein family with a cell-type-specific expression in alveolar type II cells in mice and hitherto unknown function. In type II pneumocytes, LAMP3 is localized in lamellar bodies, secretory organelles releasing pulmonary surfactant into the extracellular space to lower surface tension at the air/liquid interface. The physiological function of LAMP3, however, remains enigmatic. We generated *Lamp3* knockout mice by CRISPR/Cas9. LAMP3 deficient mice are viable with an average life span and display regular lung function under basal conditions. The levels of a major hydrophobic protein component of pulmonary surfactant, SP-C, are strongly increased in the lung of *Lamp3* knockout mice, and the lipid composition of the bronchoalveolar lavage shows mild but significant changes, resulting in alterations in surfactant functionality. In ovalbumin-induced experimental allergic asthma, the changes in lipid composition are aggravated, and LAMP3-deficient mice exert an increased airway resistance. Our data suggest a critical role of LAMP3 in the regulation of pulmonary surfactant homeostasis and normal lung function.

## Introduction

The lysosome-associated membrane glycoprotein (LAMP) family consists of five members (LAMP1, LAMP2, LAMP3 / DC-LAMP, CD68 / macrosialin, and LAMP5 (BAD-LAMP)). All family members are heavily N- and O-glycosylated type I transmembrane proteins localized to the limiting membrane of lysosomes and lysosome-related organelles (1). LAMP1 and LAMP2 are ubiquitously expressed (1), while LAMP3 (synonymously called DC-LAMP), CD68, and LAMP5 show cell-type-specific or tissue-specific expression. CD68 expression is restricted to macrophages, monocytes, and microglia (2), whereas LAMP5 is expressed exclusively in the brain (3). The LAMP3-expression pattern between humans and mice differs: LAMP3 is expressed in humans in alveolar type II (AT2) cells and dendritic cells (DC) (eponymously for its alternative name “DC-LAMP” or CD208), respectively. LAMP3 is found exclusively in AT2 cells in mice but not in DCs (4-7). These findings suggest a conserved and critical function of LAMP3 in AT2 cells. On the subcellular levels, LAMP3 is localized in lamellar bodies (LBs) (4). However, the function of LAMP3 in these cells remains enigmatic until now; if and how LAMP3 affects surfactant homeostasis, e.g., mediating surfactant release via LBs, is also poorly understood.

The primary function of AT2 cells is mainly attributed to pulmonary surfactant release to lower surface tension at the lung’s air/liquid interface. Pulmonary surfactant is a mixture of lipids and proteins (8): The major hydrophobic protein components, surfactant proteins B (SP-B) and C (SP-C), are small proteins deeply embedded into the phospholipids of the surfactant. Notably, deficiency in SP-B or SP-C due to genetic mutations in *SFTPB* or *SFTPC* cause hereditary forms of childhood interstitial lung disease (9, 10). *Sftpb* gene-targeted mice die of respiratory failure after birth, associated with irregular and dysfunctional LBs and tubular myelin (11). SP-C deficient mice have a regular life span and only show subtle deficits in lung mechanics and surfactant stability (12).

The composition of surfactant lipids has extensively been analyzed (13-17). The most abundant and characteristic lipid class is phosphatidylcholine (PC). PC makes up approximately 70% of the total surfactant mass with dipalmitoylphosphatidylcholine (DPPC; PC 16:0/16:0) as the primary lipid species (18). An overall decrease of PC lipids is coupled to higher surface tension and lower surfactant protein inclusion (8). Other hight abundant and functionally essential phospholipid classes in surfactant include phosphatidylglycerol (PG) and phosphatidylinositol (PI), while phosphatidylethanolamine (PE), and sphingomyelin (SM). Are reported as minor components of surfactant and most likely derive from other cell membranes. In addition, lysophosphatidylcholine (LPC) can be found in low abundance in pulmonary surfactant (18, 19). Preassembled surfactant is stored in secretory organelles known as LB that fuse with the cell membranes and release pulmonary surfactant. LBs are multivesicular body / late endosome-derived specialized organelles filled with surfactant characterized by a typical “myelin”-like appearance by electron microscopy. Phospholipids are directly transferred from the endoplasmic reticulum to LB through a vesicle-independent mechanism by the phospholipid transporter “ATP binding cassette subfamily A member 3” (ABCA3) (20, 21). Like *SFTPB* or *SFTPC*, loss-of-function mutations in *ABCA3* were identified in full-term infants who died from unexplained fatal respiratory distress syndrome (22). *ABCA3* mutations represents the most frequent form of congenital surfactant deficiencies (23).

A presumably loss-of-function mutation in *LAMP3* (p.(E387K)) was recently identified in an Airedale Terrier dog breed, leading to severe symptoms and pathology similar to clinical symptoms of the most severe neonatal forms of human surfactant deficiency (24). The affected puppies’ symptoms included lethal hypoxic respiratory distress and occurred within the first days or weeks of life (24). LBs were smaller, contained fewer lamellae, and occasionally revealed a disrupted common limiting membrane (24).

To conclusively address the physiological role of LAMP3 in surfactant homeostasis using a genetically defined animal model, we generated *Lamp3* knockout mice (KO, *Lamp3*^*-/-*^) by CRISPR/Cas9. In contrast to dogs bearing a natural mutation in *LAMP3, Lamp3*^*-/-*^ mice are vital, display a regular lung function at the basal level, and show no increased prenatal lethality. The lungs of *Lamp3*^*-/-*^ animals did neither macroscopically nor microscopically differ from those of wild type littermates. However, biochemical analysis of broncho-alveolar lavage (BAL) fluid clearly revealed increased pro-SP-C levels and altered lipid composition in the *Lamp3*^*-/-*^ mice. These differences are further pronounced under diseased conditions as evoked by allergen-induced experimental asthma, and most notably, diseased *Lamp3*^*-/-*^ mice revealed significantly increased airway resistance, when stressed during methacholine provocation testing. In conclusion, our data suggest a critical but not vital role of LAMP3 in pulmonary surfactant homeostasis associated with normal lung function.

## Material & Methods

### Reagents and antibodies

Analytical grade chemicals were purchased, if not stated otherwise, from Sigma-Aldrich (MO., USA). Antibodies: The following antibodies were used in the study: Rat monoclonal LAMP3, clone 1006F7.05 (Dendritics), SP-C, rabbit polyclonal (Abcam), rabbit polyclonal antibodies against mature SP-B and Pro-SP-B (Seven Hills Bioreagents) were a generous gift from Jeffrey A. Whitsett, polyclonal β-Actin (Santa Cruz Biotechnology, Inc), polyclonal rabbit antiserum against ABCA3 was a kind gift from Michael L. Fitzgerald (20). Fluorophore-conjugated secondary antibodies against the corresponding primary antibody species (AlexaFluor 488) were purchased from Invitrogen / Molecular Probes and were diluted 1:500.

### Generation and housing of Lamp3^-/-^ mice

Mice with a null mutation in the Lamp3 gene were generated in a C57BL/6N background using a CRISPR genome-editing system. For this purpose, Cas9 was combined with two single guide RNAs (sgRNAs) targeting the second exon in the Lamp3 gene with the following spacer sequences: sgRNA_Lamp3 F1: TCATCTACTGACGATACCAT and sgRNA_Lamp3 R1: GCTAGACTAGCTCTGGTTGT microinjected into fertilized zygotes. Genome editing was confirmed by PCR amplification in the resulting founder mice with the primers Lamp3F 5’-GATGGGGGAGGGATCTTTTA-3’ and Lamp3R 5’-GTTGGCCTCTGATTGGTTGT-3’. A founder harboring a 40 bp deletion within the second exon leading to a frameshift mutation was selected for further breeding. Mice were housed under specific pathogen-free conditions (12 hours’ light/dark cycle, constant room temperature, and humidity). Food and water were available ad libitum.

### Ovalbumin model of allergic asthma

Female, 6-to 8-week-old *Lamp3*^*-/-*^ and wildtype littermates mice (CAU, Kiel, Germany) were used for the allergic asthma model. They received ovalbumin (OVA)-free diet and water ad libitum. All animal studies were approved by the local animal ethics committee.

Mice were sensitized to OVA by three intraperitoneal (i.p.) injections of 10 µg of OVA (OVA grade VI, Sigma-Aldrich, St. Louis, MO, USA) adsorbed to 150 µg of aluminum hydroxide (imject alum, Thermo Fisher Scientific, Waltham, MA, USA) on days 1, 14 and 21. To induce acute allergic airway inflammation, mice were exposed three times to an OVA (OVA grade V, Sigma-Aldrich) aerosol (1% wt/vol in PBS) on days 26, 27, and 28. Control animals were sham sensitized to PBS and subsequently challenged with OVA aerosol.

### Lung function analysis

Steady-state lung parameters midexpiratory flow (EF50), frequency (f), functional residual capacity (Frc), minute volume (MV), peak expiratory flow (PEF), peak inspiratory flow (PIF), expiration time (Te), inspiration time (Ti), and tidal volume (TV) were measured in conscious, spontaneously breathing mice using FinePointe non-invasive airway mechanics double-chamber plethysmography (NAM, Data Science International, St. Paul, MN, USA). Steady-state airway resistance (RI) and dynamic compliance (Cdyn) were measured in anesthetized and ventilated mice using FinePointe RC Units (Data Science International). Airway responsiveness to methacholine (MCh, acetyl-β-methylcholine chloride; Sigma-Aldrich) challenge was invasively assessed on day 29 using FinePointe RC Units (Data Science International, St. Paul, MN, USA) by continuous measurement of RI. Therefore, animals were anesthetized with ketamine (90 mg/kg body weight; cp-pharma) and xylazine (10 mg/kg BW; cp-pharma) and tracheotomized with a cannula. Mechanical ventilation was previously described (25). Measurements were taken at baseline (PBS) and in response to inhalation of increased concentrations of aerosolized methacholine (3.125; 6.25; 12.5; 25; 50; and 100 mg/mL). After assessment of lung function, all animals were sacrificed by cervical dislocation under deep anesthesia.

### Bronchoalveolar lavage

For bronchoalveolar lavage, lungs were rinsed with 1 ml of fresh, ice-cold PBS containing protease inhibitor (Complete, Roche, Basel, Switzerland) via a tracheal cannula. Cells in BAL fluid were counted in the Neubauer chamber. Fifty microliter aliquots of lavage fluids were cytospined, and cells were microscopically differentiated according to morphologic criteria.

### Electron microscopy

Animals were anesthetized and lungs were fixed by perfusion through the right ventricle with HEPES buffer (pH 7.35) containing 1.5 % glutaraldehyde and 1.5 % formaldehyde, at a fixed positive inflation pressure of 25 cm H2O. After storage of the lungs in the same solution overnight, 3 mm sized pieces of lung tissue were postfixed first in 1 % osmium tetroxide for 2 hours and then in 4 % uranyl acetate overnight (both aqueous solutions). Samples were dehydrated in acetone and embedded in Epon. Ultratin sections of 60 nm were poststained with aqueous uranyl acetate and lead citrate (26) and imaged in a Morgagni TEM (FEI, Eindhoven, NL). All animal studies were approved by the local animal ethics committee.

### Histology

Lungs were dissected from mice, and the left lung lobe was snap-frozen in liquid nitrogen. The other lobes were fixed ex-situ with 4% (wt/vol) paraformaldehyde via the trachea under constant pressure, removed and stored in 4% paraformaldehyde. Volume of these lobes was determined based on Archimedes principle. In brief, the remaining lung lobes were placed in a water-filled container set on electronic scales. The weight of water displaced by the submerged lung equals the volume of the lung. Then lung tissues were embedded in paraffin. For analysis of lung inflammation, 2 µm sections were stained with periodic acid-Schiff (PAS). Photomicrographs were recorded by a digital camera (DP-25, Olympus, Tokyo, Japan) attached to a microscope (BX-51, Olympus) with a 20-fold magnification objective using Olympus cell^A software. Mucus quantification was performed as previously described (27). Apoptotic cells were stained using ApopTag Peroxidase In Situ Apoptosis Detection Kit (Merck Millipore, MA, USA) according to manufacturers instructions.

### Immunofluorescence staining of lung tissue

4% Paraformaldehyde-fixed lungs were incubated in cryoprotectant solution (30% w/v sucrose in 0.1 M phosphate buffer, pH 7.4) over night. 35 µm thin sections were cut with a Leica 9000s sliding microtome (Leica, Wetzlar, Germany) and collected in cold 0.1 M phosphate buffer. Free floating sections were stained by blocking in blocking solution (0.5% Triton-X 100, 4% normal goat serum in 0.1 M PB pH 7.4) for 1 hour at room temperature. Subsequently, sections were incubated in blocking solution containing the primary antibody at 4°C over night. After washing three times with wash solution (0.1 M PB pH 7.4 containing 0.25% Triton-X 100), sections were incubated for 2 hours in secondary antibody in solution, washed again three times in wash solution containing 4′,6-Diamidin-2-phenylindol (DAPI) and finally brought on glass slides and embedded in Mowiol/DABCO. Cells were imaged using a Zeiss confocal microscope equipoped with a 63x objective (Zeiss LSM980 AiryScan 2).

### Homogenization of lung tissue

Deep frozen lungs were homogenized with mortar and pestle. An aliquot of 30 mg of lung powder was transferred into RLT buffer, and RNA was isolated with RNeasy Mini Kit (Qiagen) according to the manufactures’ guidelines. 1 µg of RNA was used for cDNA synthesis (First Strand cDNA Synthesis Kit, Thermo Fisher Scientific, Waltham, MA, USA). Another 30 mg aliquot of lung powder was transferred into RIPA buffer, and proteins were isolated. Protein concentration was determined with BCA assay (Thermo Fisher Scientific, Waltham, MA, USA).

### Immunoblot analysis

Western analysis was performed on lung tissue homogenates from wildtype and LAMP3-deficient mice. Protein content was measured by the bicinchoninic acid reagent (Thermo Fisher Scientific). Defined amounts of protein were separated on SDS-PAGE and transferred to nitrocellulose membrane. Membranes were incubated with anti-pro-SP-C (rabbit monoclonal, 1:1000, Abcam, Cambridge, UK), anti-pro-SP-B (rabbit polyclonal, 1:2000, Seven Hills Bioreagents, CA, USA), anti-mature SP-B (rabbit polyclonal, 1:1000, Seven Hills Bioreagents), both generous gifts from Jeffrey A. Whitsett, and anti-β-actin (mouse monoclonal, 1:200, Santa Cruz Biotechnology, Heidelberg, Germany). Donkey anti-rabbit IgG HRP (1:2000, Santa Cruz) and donkey anti-mouse IgG HRP (1:2000, Cell Signaling) served as secondary Abs. Immunoreactive proteins were visualized using the ECL Western blotting detection system (Bio-Rad Laboratories GmbH, Feldkirchen, Germany), band intensity was quantified with Image J 1.52 (NIH), and data were normalized to β-actin levels.

### qPCR

Quantitative RT-PCR was performed on the Roche LightCycler 480 Instrument II system using the LightCycler 480 SYBR Green I Master (Roche Applied Science, Mannheim, Germany). Template cDNA was diluted 1:10. RT-PCR was performed in triplicates in a total volume of 10 µl according to the manufacturers’ instruction with a final primer concentration of 0,5 µM. The following primer sequences for housekeeping gene RPL32 were used: forward 5‘-AAAATTAAGCGAAACTGGCG-3’, reverse 5‘-ATTGTGGACCAGGAACTTGC-3’. QuantiTect Primer Assays from Qiagen (Venlo, The Netherlands) were used for *Sftpb* (QT00124908) and *Sftpc* (QT00109424). Negative controls were included to detect possible contaminations. Amplification specificity was checked using the melting curve and agarose gel electrophoreses. Specific mRNA levels were normalized to the level of the housekeeping gene RPL32 in the same sample.

### Chemicals and lipid standards for lipidomics

All solvents, reagents, and lipid standards were used in the highest available purity. Internal standards were acquired from Avanti Polar Lipids (Avanti Polar Lipids, Alabama, USA). Storage solution was created by combining chloroform, methanol, and water (20/10/1.5; v/v/v. ESI spray mixture was created by combining chloroform, methanol with added 0.1% (wt/v) ammonia acetate and 2-propanol (1:2:4; v/v/v).

### Lipid extraction

A customized methyl-tert-butyl ether (MTBE)-based lipid extraction method (28) was applied for BAL and lung tissue homogenates. Details can be found in the **Supplementary Methods**. Lipid extracts were stored in a storage solution at -20 °C until further usage. Cholesterol determination was performed as described earlier (29) (**Supplementary Methods**).

### Shotgun lipidomics measurements

Aliquots of 10 µL of the lipid extract and cholesterol derivatization were diluted in 190 µL ESI spray mixture for each measurement, vigorously mixed, and centrifuged prior to loading into a 96 well plate (Eppendorf, Hamburg, Germany). The well plate was sealed with aluminum sealing foil and kept at 15 °C during the measurement process. All measurements were performed in duplicates. All mass spectrometric acquisitions were performed using a Triversa Nanomate (Advion, Ithaca, USA) as an autosampler and nano-ESI source, applying a spray voltage of 1.1 kV and backpressure of 1.1 psi in ionization modes. For the acquisition of tandem mass spectrometric data, a Q Exactive Plus was used (Thermo Fisher Scientific, Bremen, Germany; methodical details are found in the **Supplementary Methods**).

### Lipidomics data interpretation

For data interpretation, the software pipeline described earlier was utilized (see details **Supplementary Methods**) (30). Briefly, mass spectra raw files were converted to mzML format with the software module “msconvert” from ProteoWizard version 3.0.18212-6dd99b0f6 (31). Converted files were then imported into LipidXplorer version 1.2.8.1 (32). For each ESI mode, a different set of MFQL (33) files is used. The files generated in LipidXplorer were used in lxPostman for further post-processing, including quality control and quantitation.

### Statistical analysis for lipidomic data

Technical replicates are averaged and normalized on protein content. Missing data for ANOVA and PCA were imputed via MetImp 1.2 (34) MCAR/MAR mode with Random Forrest was used as a method with a group-wise missing filter set to 70%. Data were analyzed via Cluster 3.0 version 1.58 and Java Treeview version 1.1.6r4 (35), R version 3.6.1 and RStudio version 1.2.1335 and GraphPad Prism version 8.4.2, and Microsoft Excel for Microsoft 365. Two-way ANOVA was performed with the parameters matching time points for each row, mixed-effects with full fitting, and within each row comparing columns with correcting for multiple comparisons via Tukey. Lipid values are expressed as mean ± standard deviation and significance is designated as *, P < 0.05; **, P < 0.01; ***, P < 0.001. For easier visualization of significant lipid results, lipids from two-way ANOVA were autoscaled. In this procedure, data is centered on the means of groups and then divided by the standard deviation, removing the impact of differences between lipid abundance.

### Bubble Surfactometer

Cell free BAL was weighed (=400-800 µL) and ultra-centrifuged (J2-MC, Beckman Coulter, Krefeld, Germany) for 1h at 4°C at 38.730 g. The supernatant was decanted. The pellet was weighed and resuspended using saline containing 1.5 mmol/L CaCL_2_ equivalent to a concentration of 1:50 (w/w) compared to the basic BAL. Two µL of the concentrated BAL were analyzed for content of choline-containing phospholipids (DPPC, lysolecithin, sphingomyelin; LabAssay™ Phospholipids, FUJIFILM WAKO). Then, samples were normalized to final phospholipids-, i.e. choline containing phospholipids-, concentrations of 1.5 mg/ml and analyzed in the Pulsating Bubble Surfactometer (PBS; (36)). A PBS-sample chamber was prefilled using degassed 10% sucrose containing saline plus 1.5 mmol/L CaCL_2_. Then, 5 µL of each normalized BAL sample were pipetted on the top of the sucrose prefill close to the chimney, where it remains by buoyancy. Subsequently, to mounting of the chamber in the PBS, an air bubble is created by aspiration of ambient air through the chimney and phospholipids might adsorb 30s to the air liquid interface. Then, the bubble is cyclically expanded and compressed at a frequency of 20/min, a compression rate of 50% surface area and at 37° C. Surface tension at maximum (gmax) and minimum (gmin) bubble size is calculated by the PBS and recorded for a 300s period.

### Statistical analyses

If not stated otherwise, a two-tailed unpaired t-test was performed using GraphPad Prism Software Version 5.03. Significant values were considered at P < 0.05. Values are expressed as mean ± standard error of the mean (SEM) and significance is designated as *, P < 0.05; **, P < 0.01; ***, P < 0.001.

## Results

### *Lamp3* knockout mice are viable with normal lung anatomy and pulmonary function

LAMP3 is expressed highest in humans and mice in the lung’s AT2 cells, where it localizes to LBs (4, 5). Notably, a naturally occurring recessive missense *LAMP3* variant has recently been associated with a fatal neonatal interstitial lung disease in Airedale Terrier dogs (24). Therefore, we aimed to analyze the function of LAMP3 in the lung in a genetically more defined animal model and generated *Lamp3* knockout mice (*Lamp3*^*-/-*^). CRISPR/Cas9 editing of the Lamp3 allele in mice with guide RNAs targeting exon 2 (**Figure 1A, B**) yielded a 40 bp deletion resulting in an early Stop-codon in one founder and the mutation was transmitted in the germline (**Figure 1C**). PCR with primers spanning this deletion was used for genotyping (**Figure 1D**). LAMP3 was readily detectable by immunohistochemistry on lung sections with a monoclonal antibody in AT2 cells of wildtype mice but was completely absent in *Lamp3*^*-/-*^ mice, validating the knockout at the protein level (**Figure 1E**). We hypothesized that knockout of *Lamp3* in mice causes abnormal surfactant homeostasis and postnatal death. However, in contrast to dogs with a *LAMP3* mutation, homozygous *Lamp3*^*-/-*^ mice were surprisingly born according to the expected Mendelian distribution and survived the weaning phase without increased mortality (**Figure 1F**). No premature death was observed until 15 months-of-age, indicating that LAMP3 is not essential for survival in mice. Macroscopically, *Lamp3*^*-/-*^ mice were indistinguishable from wildtype littermates with no obvious phenotype. Histology and analysis of semi-thin sections revealed normal micro-anatomy of the lung with a typical distribution of AT2 cells, type I pneumocytes, alveolar macrophages, and regular capillaries (**Figure 1G**). Macroscopic signs of multifocal emphysema or edema (as previously described in dogs with a mutation in *LAMP3*, (24)) were absent in *Lamp3*^*-/-*^ mice (**Figure 3H**). The lung volume, normalized to the body weight in 3-months-old animals, was comparable between wildtype and *Lamp3*^*-/-*^ mice (**Figure 1I**).

**Figure 1.**
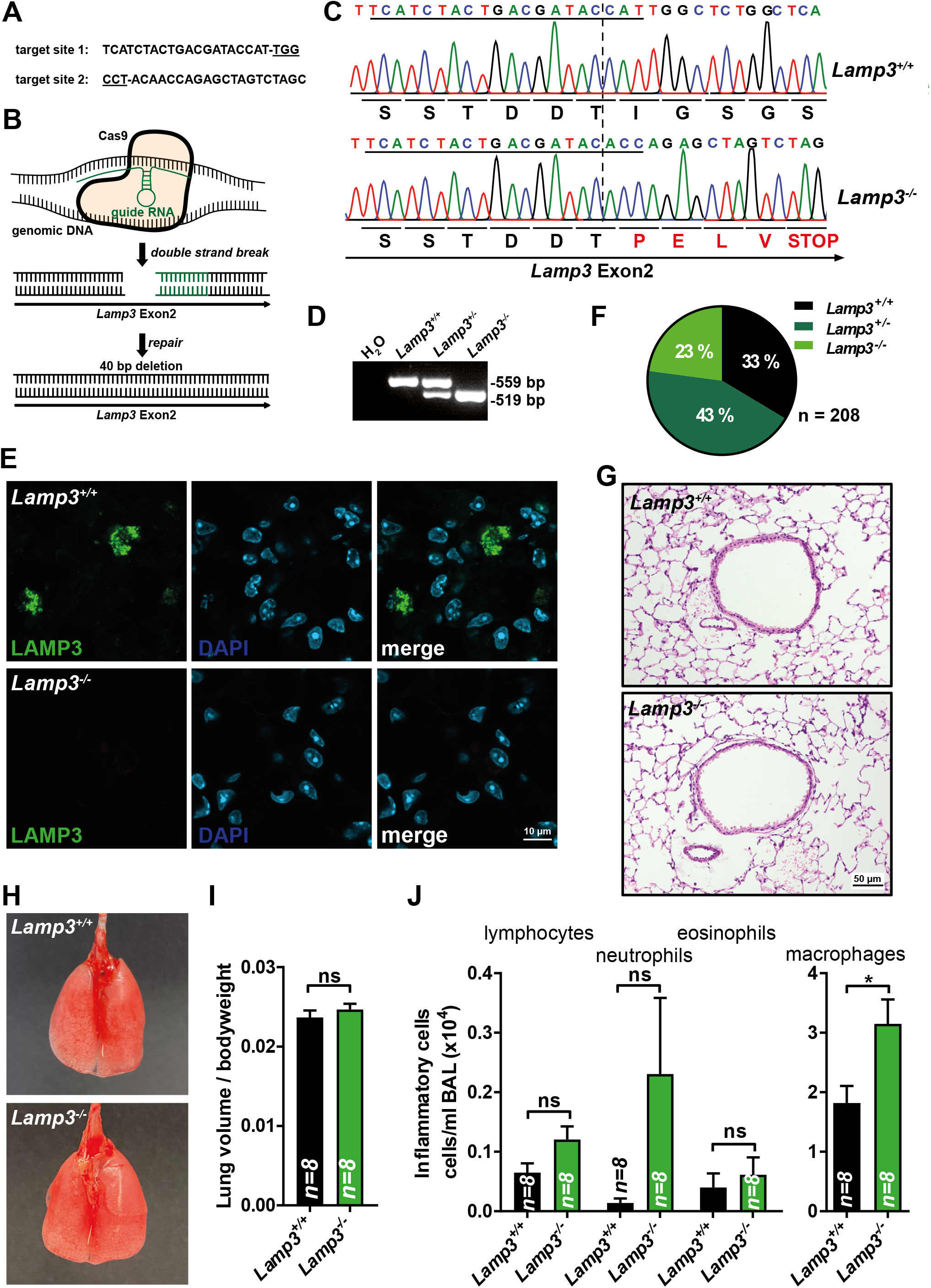
CRISPR/Cas9-mediated knockout of *Lamp3* in mice. **(A)** CRISPR guide-RNA sequences used for targeted editing of the murine *Lamp3* allele. The Protospacer Adjacent Motif (PAM) is underlined. **(B)** Schematic drawing of CRISPR/Cas9-mediated targeting of exon 2 of the *Lamp3* locus. **(C)** Sanger sequencing chromatogram of a PCR-product covering the targeted segment with genomic tail-DNA of one wild type and one homozygous *Lamp3*^*-/-*^ mouse as a template. CRISPR target site one is underlined. The frameshift resulting from endogenous repair mechanisms results in an early stop codon. **(D)** 2% Agarose gel with PCR-products of a PCR covering parts of exon 2 of the *Lamp3* locus used for genotyping. **(E)** Immunohistochemistry staining of lung sections of wildtype and *Lamp3*^*-/-*^ mice with an antibody specific for LAMP3. **(F)** Genotype distribution of the litters from heterozygote *Lamp3*^*+/-*^-breeding pairs after weaning (3-4 weeks after birth). N = 208 individuals. **(G)** Hematoxylin / Eosin staining of lung sections from adult wildtype and *Lamp3*^*-/-*^ mice. **(H)** Photos of the inflated lungs of adult wildtype and *Lamp3*^*-/-*^ mice. **(I)** Lung volume/body weight ratio of wild type and *Lamp3*^*-/-*^ mice. n = 8 (per genotype), ns = not significant. **(J)** The number of inflammatory cells (lymphocytes, neutrophils, eosinophils, and macrophages) in the BAL of wildtype and *Lamp3*^*-/-*^ mice. n = 8 (per genotype), ns = not significant; 0.05; * = p ≤ 0.05.

**Figure 2.**
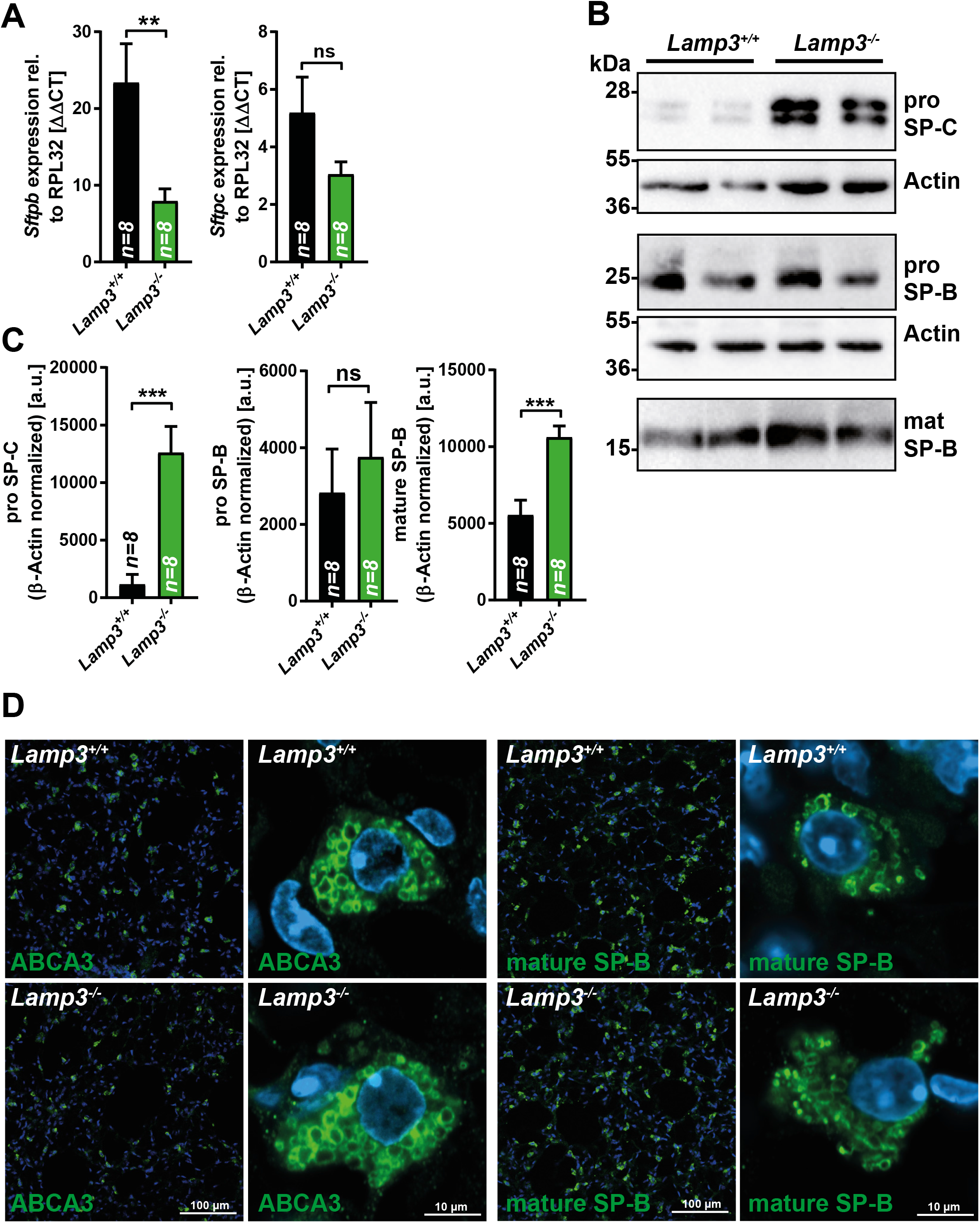
LAMP3-deficiency results in altered (pro-)surfactant C levels. **(A)** Quantitative real-time PCR of total RNA from wildtype and *Lamp3*^*-/-*^ mice for SP-B and SP-C. The expression is normalized to the housekeeping gene RPL32. n = 8 (per genotype), ns = not significant; ** = p ≤ 0.01. **(B)** Representative immunoblots of lung tissue lysates from wildtype and *Lamp3*^*-/-*^ mice for pro-SP-C, pro-SP-B, and mature SP-B. Actin is depicted as a loading control. **(C)** Quantification of immunoblots for pro-SP-C, pro-SP-B, and mature SP-B normalized to Actin as a loading control. n = 8 (per genotype); ns = not significant; *** = p ≤ 0.001. **(D)** Immunofluorescence staining of lung sections from adult wildtype and *Lamp3*^*-/-*^mice for ABCA3 and mature SP-B (green). Nuclei are stained with DAPI (blue).

**Figure 3.**
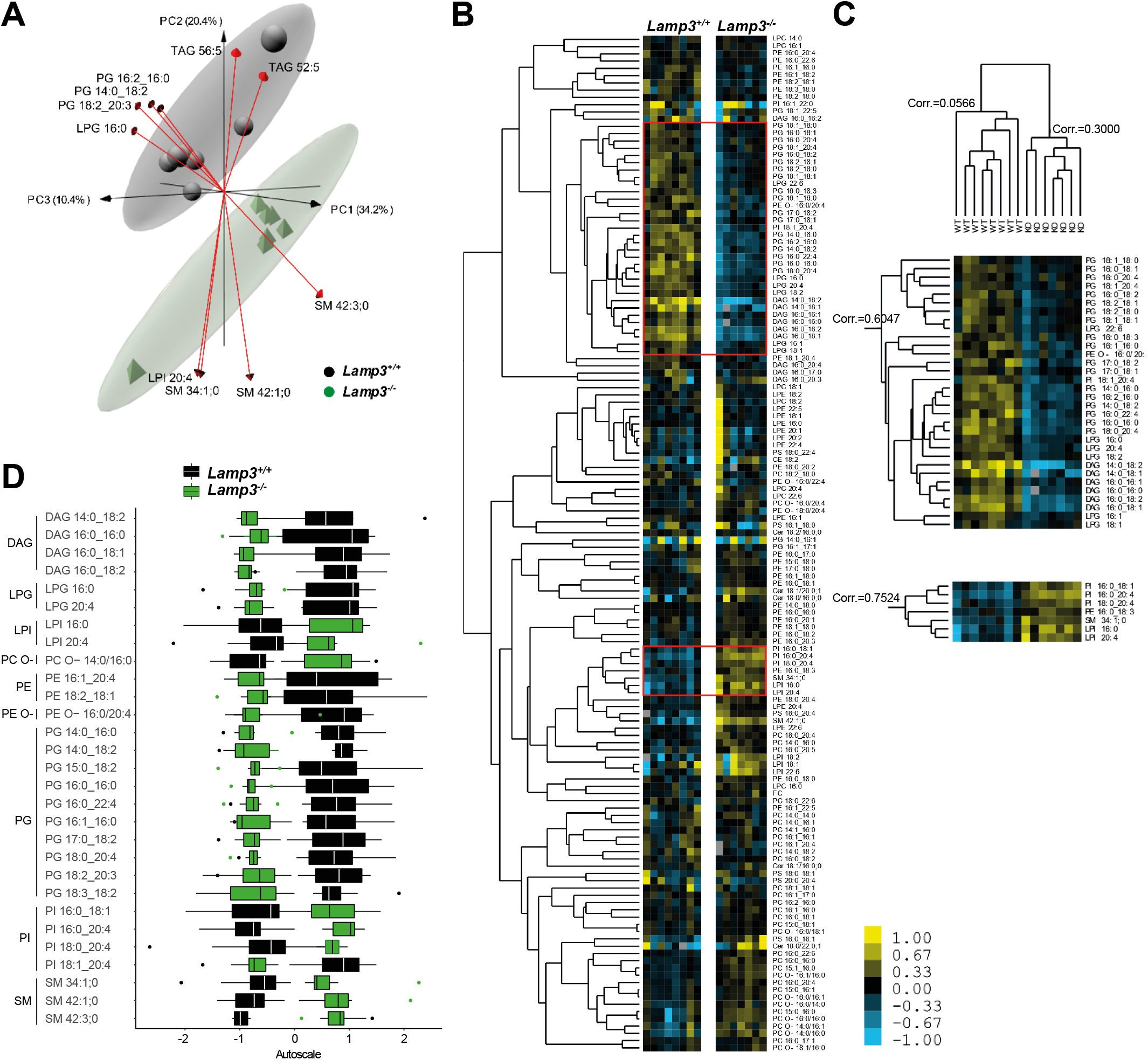
Deficiency of LAMP3 results in minor changes in the BAL lipidome under basal conditions. **(A)** Three-dimensional principal component analysis (PCA) loadings plot of the BAL lipidome of wild type (black) and *Lamp3*^*-/-*^ (green) mice. Each point represents one individual animal. Arrows show the 10 lipid species with the strongest effect across all principal components. Ellipses show the 95% confidence interval of the group. PC1 to PC3 explain 65% of the variability in the data set. **(B)** Hierarchical clustering of 140 lipid species (of 208 after application of a 90% occupation threshold) identified in eight wild type and seven *Lamp3*^*-/-*^ mice BAL fluid samples. Each row represents a lipid species and each column a sample. Red boxes indicate enlarged selections **(C2)** and **(C3). (C1)** The hierarchical clustered tree of sample groups correlates both groups into two major branches. **(C2)** The Upper selected segment of the HCA with 31 lipids shown with a relative decrease to wildtype mice. **(C3)** The lower selected segment of the HCA was shown with a relative increase of 7 lipids to the control. **(D)** Differences in the BAL lipidome of wild type and *Lamp3*^*-/-*^ mice mutant using autoscaled ranges. Samples are color-coded according to their group, black for *Lamp3*^*+/*+^ and green for *Lamp3*^*-/-*^ mice. Boxplots show the median with the interquartile range. Whiskers show maximum and minimum with outliers in the respective colors. Data is on the basis of mole percentage. Only lipid species are shown with a significant difference computed with a two-way ANOVA.

Furthermore, lung function analysis revealed no abnormalities in the breathing pattern of *Lamp3*^*-/-*^ mice regarding the frequency, flow, volume, and time parameters as non-invasively assessed in spontaneously breathing animals as well as for airway resistance and compliance as recorded invasively in relaxed, ventilated animals (**Table 1**). Analysis of the BAL cellular composition revealed a tendency towards a higher number of lymphocytes and neutrophils in *Lamp3*^*-/-*^ mice, which, however, did not reach statistical significance. The number of macrophages was moderately increased in *Lamp3*^*-/-*^ mice (**Figure 1J**).

**Table 1.** Steady state lung parameters in wildtype and *Lamp3*^*-/-*^ mice. Airway resistance (RI) and dynamic compliance were measured in anesthetized and ventilated mice using FinePointe RC Units. Midexpiratory flow (EF50), frequency (f), functional residual capacity (Frc), minute volume (MV), peak expiratory flow (PEF), peak inspiratory flow (PIF), expiration time (Te), inspiration time (Ti), and tidal volume (TV) were measured in conscious, spontaneously breathing mice using FinePointe non-invasive airway mechanics (NAM) double chamber plethysmography. Values displayed as mean ± SEM, n= 7 per genotype.

| parameter (abbr.) | unit | mean ± SEM (WT) | mean ± SEM (KO) | p-value |
| --- | --- | --- | --- | --- |
| airway resistance (RI) | cmH<sub>2</sub>O*secs/mL | 1.423 ± 0.19 | 1.282 ± 0.16 | 0.5869 |
| dynamic compliance (Cdyn) | mL/cm H<sub>2</sub>O | 0.041 ± 0.01 | 0.051 ± 0.01 | 0.5126 |
| midexpiratory flow (EF50) | ml/sec | 5.028 ± 0.32 | 4.958 ± 0.31 | 0.8772 |
| frequency (f) | BPM | 235.6 ± 8.1 | 234.0 ± 9.2 | 0.8968 |
| functional residual capacity (Frc) | ml | 0.7667 ± 0.14 | 0.7275 ± 0.05 | 0.7946 |
| minute volume (MV) | ml/min | 123.9 ± 7.9 | 121.2 ± 6.7 | 0.7985 |
| peak expiratory flow (PEF) | ml/sec | 5.995 ± 0.38 | 5.884 ± 0.30 | 0.8215 |
| peak inspiratory flow (PIF) | ml/sec | 5.571 ± 0.38 | 5.314 ± 0.31 | 0.6088 |
| expiration time (Te) | sec | 0.1308 ± 0.005 | 0.1308 ± 0.005 | 0.9982 |
| inspiration time (Ti) | sec | 0.1281 ± 0.004 | 0.1300 ± 0.004 | 0.7522 |
| tidal volume (TV) | ml | 0.5234 ± 0.018 | 0.5190 ± 0.014 | 0.8480 |

### LAMP3 deficiency in mice causes changes in surfactant protein levels

We reasoned that a lack of LAMP3 might affect the levels of the surfactant proteins SP-B and SP-C, major components of the LB-stored- and secreted surfactant. Indeed, quantitative PCR analysis revealed reduced levels of *Sftpb* transcript and a similar trend (though not statistically significant) for *Sftpc* in total lung RNA (**Figure 2A**) of *Lamp3* knockout mice. In contrast, immunoblot analysis of lung tissue lysates from wildtype and *Lamp3*^*-/-*^ mice for SP-B and SP-C showed normal levels of pro-SP-B and a striking increase in the levels of (pro-) SP-C, indicating a critical role of LAMP3 in the biosynthesis or turnover of SP-C (**Figure 2B, C**). The levels of mature SP-B were also significantly increased, though only to a moderate level compared the increase in SP-C.

Immunofluorescence analysis on lung sections from wildtype and *Lamp3*^*-/-*^ mice revealed no major differences in the staining pattern of the essential LB proteins ABCA3 or mature SP-B. Both proteins localized in wildtype and *Lamp3*^*-/-*^ mice to ring-like structures of AT2 cells, representing LBs as determined by high-resolution fluorescence microscopy (**Figure 2D**).

### The lipid composition of the bronchoalveolar lavage is altered in *Lamp3*^*-/-*^mice

Given the apparent differences in SP-C levels and mature SP-B in protein extracts of lung tissue lysates, we next analyzed the lipid composition of the BAL and total lung homogenates by a semi-targeted shotgun lipidomics approach. After shotgun lipidomics, a total of 208 different lipid species in the BAL were quantified. We first determined if the two groups (wildtype and *Lamp3*^*-/-*^ BAL lipids) could be separated by analyzing the lipidomics dataset by unsupervised principal component analysis (PCA) (**Figure 3A**). The apparent separation was characterized by increased PGs and LPG levels in the wildtype while SMs are increased in the *Lamp3*^*-/-*^ mice. To gain better insight into the perturbations on the lipid species level, we further applied hierarchical clustering (**Figure 3B)**. In conjunction with the PCA, wildtype and *Lamp3*^-/-^ mouse BAL lipid profiles were clearly distinguished by the underlying Spearman Rank correlation. According to the genotype, the abundance of lipid species’ correlated for PG, LPG, and DAG, showing a relative increase in wildtype mice, which was also confirmed by statistical analysis for the selected indicated lipids species (**Figure 3C)**. In contrast, PI and LPI are decreased in *Lamp3*^-/-^ mice. Notably, PG is an essential component of the pulmonary surfactant, and as multiple lipids of the same class are affected, a disruption of the normal BAL physical properties is likely. In contrast to BAL’s lipid profiles, no statistical differences were identified between wild type and *Lamp3*^*-/-*^ mice by lipidomic analysis of the perfused total lung tissue (**Supplemental Figure 1A-C**). These findings, in summary, indicate that LAMP3 deficiency leads to subtle but statistically significant changes in the lipidome and particularly surfactant-relevant lipids of the BAL.

### *Lamp3* knockout leads to changes in the ultrastructure of lamellar bodies and altered biophysical properties of pulmonary surfactant

Defects in surfactant homeostasis are typically accompanied by a change of the ultrastructure of LBs (11, 12, 24). This finding prompted the analysis of the *Lamp3*^-/-^ lungs by transmission electron microscopy, focusing on LBs in the AT2 cells. Lamellar bodies from wildtype mice had the typical lamellar appearance (**Figure 4A**). In *Lamp3*^-/-^ mice, conspicuous LBs could occasionally be observed within AT2 cells (**Figure 4A**). They consisted of several clews of lipid lamellae and seemingly homogenous areas that were surrounded by a common limiting membrane. At higher magnification, the homogenous areas also revealed lamellae that were not immediately obvious at lower magnification (**Figure 4A**). Accordingly, we tested the surfactant’s biophysical properties and functionality obtained from BAL of wildtype and *Lamp3*^*-/-*^ mice to address any functional relevance of such morphological alterations. Surfactant samples were investigated during dynamic compression of an air bubble in a pulsating bubble (36). The surface tension of wildtype and *Lamp3*^*-/-*^ mice is summarized at minimum (gmin) (**Figure 4B**) and maximum (gmax) bubble size (**Figure 4C**). Surfactant standardized at a phospholipid concentration of 1.5 mg/ml of both wildtype and *Lamp3*^*-/-*^ mice reach a low physiological gmin close to 0 mN/m during the observational period of 300s, respectively (**Figure 4B**). During the compression period, gmin (0.95 vs 3.6 mN/m, median, p<0.01) and gmax (29.2 vs. 29.9 mN/m, median, p<0.0001) in *Lamp3*^*-/-*^ mice were found slightly but significantly lower compared to wild type surfactant. Thus initially, gmin of samples from *Lamp3*^*-/-*^ mice decreases faster, reaching <10 mN/m already after 40s compared to wildtype surfactant after 80s (mean) (**Figure 4C**), indicating a slightly increased adsorption property of surfactant from *Lamp3*^*-/-*^ mice.

**Figure 4.**
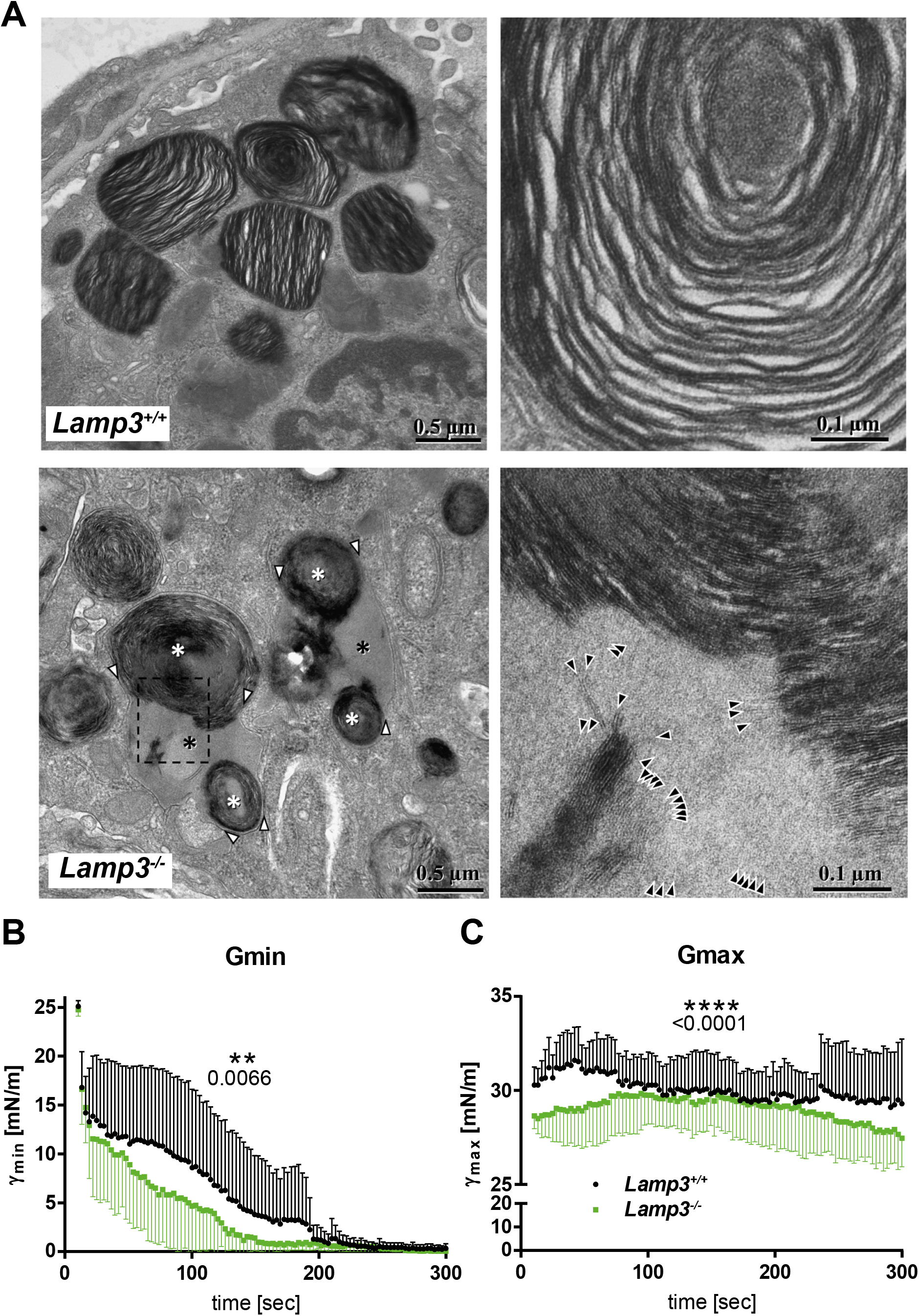
Deficiency of LAMP3 leads to altered LB morphology and reduced surfactant functionality, **(A)** Transmission electron microscopy of LB in an alveolar AT2 cells of a wildtype and a *Lamp3*^*-/-*^ mouse. Wildtype: Individual Lamellar Bodies are clearly discernible and separated (left), containing continuous lipid lamellae (right). *Lamp3*^*-/-*^: Left: Multiple clews of lipid lamellae (white asterisks) are located within shared limiting membranes (white arrowheads). The space between the clews (black asterisks) is entirely filled with light grey material. Right: enlargement of the boxed area indicated left. Higher magnification reveals lamellae also inside the light grey material, which in some cases obviously are continuations of the darker stained lamellae inside the clew (top) or the stack of lamellae bottom left. **(B, C)** Surface tension at minimum (gmin; **C**)) and maximum (gmax; **B**) bubble size, derived from a pulsating bubble surfactometer (36) during a compression period of 300s. Eight pellets of BAL from *Lamp3*^*-/-*^ and wild type mice were prepared by ultracentrifugation and resuspended in saline CaCl_2_ to standardize phospholipids at 1.5 mg/ml each. Phospholipid concentration was determined using a commercial kit. Data are plotted as mean +/-SD. Statistical analysis was performed using an unpaired t-test.

### *Lamp3*^*-/-*^ mice show increased airway resistance and aggravated BAL lipid changes under pathological conditions

The overall phenotype of *Lamp3*^-/-^ mice was mild and did not affect lung function to the extent that functional lung tests could reveal alterations in healthy, spontaneously breathing animals (**Table 1**). We, therefore, wondered if LAMP3 is more critical under challenged, i.e., pathological conditions. We applied a well-described model of experimental allergic asthma (25, 37) (**Figure 5A**) that is characterized by allergic inflammation of the airways, mucus hyperproduction, and the development of airway hyperresponsiveness (AHR), thus, resembles the pathophysiologic hallmarks underlying the symptoms of human asthma. Analysis of the different immune-cell populations (macrophages, lymphocytes, neutrophils, eosinophils) infiltrating the bronchoalveolar lumen in diseased animals did not unveil any differences between wildtype and *Lamp3*^*-/-*^ mice. The number of mucus-producing goblet cells and the amount of mucus stored in the airway mucosa did also not differ (**Supplemental Figure 3**). The degree of apoptotic cell death as revealed by terminal deoxynucleotidyl transferase dUTP nick end labeling (TUNEL) staining between wildtype and *Lamp3*^*-/-*^ mice remained unchanged (**Supplemental Figure 4**). Importantly, when animals were subjected to MCh-provocation testing to determine the degree of AHR, *Lamp3*^*-/-*^ mice displayed significantly increased airway resistance in response to MCh-inhalation (**Figure 5B**).

**Figure 5.**
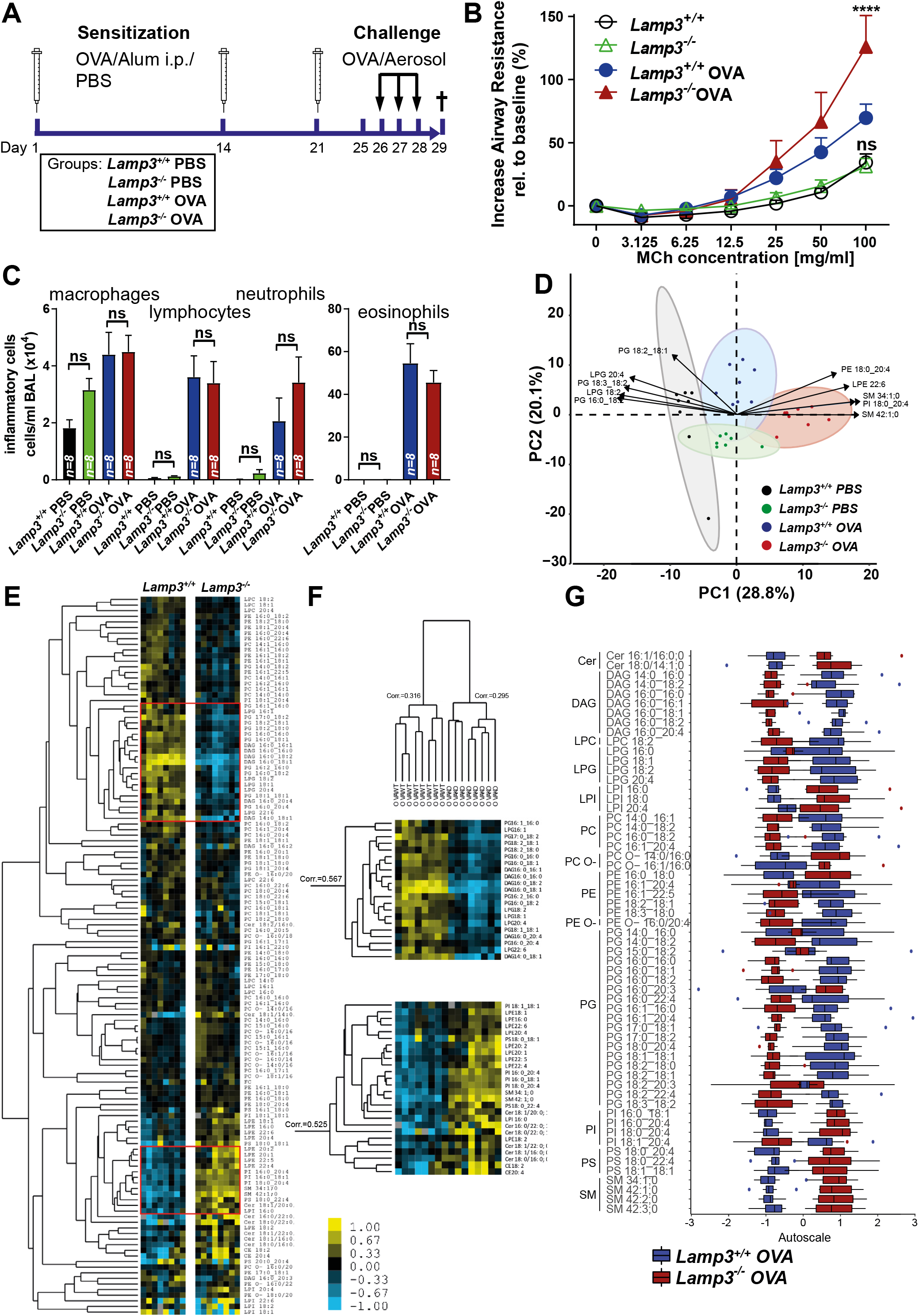
The stress-induced allergic asthma model aggravates phenotypes in *Lamp3*^*-/-*^ mice and causes a genotype-dependent increase in airway resistance. **(A)** Experimental setup of the stress-challenged model of allergic asthma and the experimental groups. **(B)** Airway resistance of the four experimental groups (wildtype, *Lamp3*^*-/-*^, wildtype + OVA, *Lamp3*^*-/-*^ +OVA). N = 8 (per genotype), significances calculated for methacholine provocation test after 100 mg/ml methacholine exposure, ns = not significant; 0.05; **** = p ≤ 0.0001 **(C)** Number of inflammatory cells in the BAL of wildtype, *Lamp3*^*-/-*^, wildtype + OVA, *Lamp3*^*-/-*^+OVA mice. N = 8 (per genotype), ns = not significant; 0.05; * = p ≤ 0.05. **(D)** Two-dimensional PCA loadings plot of the BAL lipidome of wild type (black), *Lamp3*^*-/-*^ (green), wild type +OVA (blue), and *Lamp3*^*-/-*^ +OVA (red) mice. The datasets for wild type (black), *Lamp3*^*-/-*^ (green) are identical to (**Figure 3A)**. Each point represents one individual. Arrows show the 10 lipid species with the strongest effect on all principal components. Ellipses indicate the 95% confidence interval of the experimental groups, where PC1 and PC2 describe 48.9% of the variability in the data set. The separation of the groups is in PC1, and PC2 is not complete for all groups. A clear separation between the *Lamp3*^*+/+*^ +PBS and *Lamp3*^*+/+*^ +OVA can be seen. The separation between untreated *Lamp3*^*-/-*^ and treated *Lamp3*^*-/-*^ is not complete. **(E)** Hierarchical clustering of 140 lipid species (of 208 after application of a 90% occupation threshold) identified in 8 wild type +OVA and 8 *Lamp3*^*-/-*^ +OVA mice BAL fluid samples. Each row represents a lipid species and each column a sample. Red boxes indicate selections shown in **(F)**. The two major branches of the tree represent the genotypes without false assignments. Selected segments of the hierarchical clustered tree where a relative decrease for 21 lipids for *Lamp3*^*-/-*^, mainly DAG and PG species, compared to the *Lamp3*^*+/+*^ mice are observed. The lower selection shows, in contrast, a cluster of 26 lipids with a relative increase in quantity in the mutant compared to *Lamp3*^*+/+*^. **(G)** Differences in the BAL lipidome of wild type +OVA and *Lamp3*^*-/-*^ +OVA applying autoscaling. Samples are color-coded according to their group, red for *Lamp3*^*+/+*^ +OVA and blue for *Lamp3*^*-/-*^ +OVA. Boxplots show the median with the interquartile range, and whiskers show maximum and minimum with outliers in the respective colors. Data is based on mole percentage. Selected lipid species showed a significant difference by two-way ANOVA. **(H)** Bar plots of absolute lipid values normalized on protein content. Error bars signify standard deviations.

To further investigate the impact of experimental asthma in mice, the lipid composition of the BAL and total lung tissue was determined (**Figure 5D-G**). The response to experimentally induced asthma led to further divergence in the BAL lipid composition between *Lamp3*^*-/-*^ mice and wildtype. Further increase in PG, LPG, and DAG abundance was observed in wildtype mice while levels of SM and PI increased in in *Lamp3*^*-/-*^. In the lipidomes of the perfused total lung tissue, no further changes were observed (**Supplemental Figure 2**). Hierarchical cluster analysis further revealed that the opposite response in the BAL lipidome of the wildtype and *Lamp3*^*-/-*^ mice affected a wider range of lipid species, compared to the healthy animals (**Figure 5E, Figure 3B-D**). Spearman Rank correlation between the two distinct groups (wildtype and *Lamp3*^*-/-*^) further confirmed this lipid phenotype by high correlation factor for the two major branches comprising lipid compositions of 8 animals in each group (**Figure 5F**). Lipid species of mainly PG, LPG, and DAG classes show a relative increase in *Lamp3*^*+/+*^ mice, which were statistically significant for indicated lipids species. In contrast, PI, LPE, Cer, and SM showed increased abundance in *Lamp3*^*-/-*^ mice (**Figure 5G**). In summary, the lipid homeostasis for surfactant production in BAL of *Lamp3*^*-/-*^ mice are more disturbed under the pathophysiologic conditions of experimental asthma when compared to the unchallenged animals.

## Discussion

The two best-characterized family members of the LAMP family, LAMP1 and LAMP2, are ubiquitously expressed and quantitatively account for a substantial fraction of the lysosomal membrane proteome (21, 38). Though the function of LAMP1 and LAMP2 are still not fully understood, they are supposed to play pivotal roles in the fusion of membranes, autophagy, microtubule-dependent positioning of lysosomes, and especially as structural components of the lysosomal membrane (39). In contrast, the expression of the structurally related LAMP3 protein is limited to specific cell types, implicating a more defined and tissue/cell-type-specific function. Together with the fact that LAMP3 localizes to LBs rather than lysosomes, a critical function of LAMP3 in LB function and surfactant homeostasis is suggested. In fact, the recent finding that a natural mutation (p.(E387K) in *LAMP3* leads to clinical symptoms and pathology similar to the most severe neonatal forms of surfactant deficiency in an Airedale Terrier dog bred suggested a critical role in surfactant biology and lung physiology for LAMP3.

Therefore, we aimed to investigate the role of LAMP3 in lung physiology by using a genuine knockout model. To our surprise, our findings obtained from *Lamp3*^*-/-*^ mice revealed striking differences to those found in dogs with the recessive missense LAMP3 variant: While lethal hypoxic respiratory distress and failure lead to the premature death of dog puppies within the first days or week of living, *Lamp3-*^*-/-*^ mice developed normally and did not display signs of lung pathology and dysfunction after birth. We can only speculate about the apparent difference between the different animal models: On the one hand, in contrast to laboratory mouse strains, dog breeds like the Airedale Terrier were generated by inbreeding that aimed to pronounce or produce distinct anatomical features. Thus, it could not be excluded that other genetic factors, possibly homozygous in the inbred dogs, impact this phenotype. On the other hand, the point mutation (p.(E387K)) found in the dogs resulted in the expression of a LAMP3 variant that could still be detected with a LAMP3-specific antibody in lung tissue, suggesting that the overall protein structure of the LAMP3 variant is at least in part untouched by the mutation. This finding leaves the possibility of an impaired function of the mutated variant, which is not the case when *LAMP3* is completely knocked-out by CRISPR/Cas9. These findings, however, should be considered when human patients suffering from surfactant deficiencies are analyzed: Our data suggest that *LAMP3* mutations might also present with a mild clinical phenotype, maybe not immediately presented with the typical signs of a full surfactant deficiency in human patients suffering from childhood interstitial lung disease. Since neither snap-frozen tissue samples nor BAL from the affected Airedale Terrier puppies could be obtained, characterization of the phenotype caused by a mutation of the LAMP3 gene in dogs is quite limited in regard to the lipidome profile of BAL and the function of the surfactant.

The lipidomics analyses performed in this study are in good agreement with earlier reported quantities of main lipid species and classes by Prueitt et al. (16). We find PC at 67.1 mol% in wildtype mice (16 mol% neutral lipids excluded as in study) compared to the previously reported 75.2 mol% (16). The main contributing PC class lipid species is DPPC (PC 16:0/16:0) with 43.9% compared to the 30-60% reported in the literature (14, 17, 40, 41). PG was found at 9 mol% in agreement with the values reported by others (16). Our analyses show a clear separation of the genotypes according to different lipid compositions and upon induction of experimental asthma. In the lipidomes of the perfused lung tissues, only minor *Lamp3* genotype-dependent changes were observed, which underlines the importance of LAMP3 to regulate the lipid secretion in the surfactant. It is further noteworthy that the BAL lipid composition was mostly affected by the reduction in the abundance of PGs, LPG, and DAG in the *Lamp3*^*-/-*^ mice.

Importantly, experimentally induced asthma further reduced the levels of essential surfactant lipids and in parallel, led to an increase in lipids like SM, PI, and PS. Interestingly lipids containing arachidonic acid (AA, FA 20:4) were elevated in *Lamp3*^*-/-*^ mice. Further affects by signaling was not investigated in this study, where AA plays a crusial role as educt of numerous lipid mediators. These lipids might further indicate a higher degree of cell lysis in conjunction with increased number of immune cells in BAL after OVA treatment and an excessive immune reaction. However, it should be noted that we did not observe major differences in the number of apoptotic cells in lung sections.

While our data support a role of LAMP3 in surfactant homeostasis, we can only speculate about the mechanisms of how LAMP3 might regulate surfactant levels secreted by AT2 cells. Different scenarios are feasible: On one hand, it has been speculated previously that LAMP3 plays a role in surfactant recycling by retrieving remnants of secreted LB and re-endocytosis (4). On the other hand, other LAMP family members are acting as accessory subunits of multi-transmembrane spanning proteins like LAMP1/LAMP2 for the lysosomal polypeptide transporter TAPL (42). Likewise, UNC-46, the *C. elegans* orthologue of LAMP5, acts as a trafficking chaperone, essential for the correct targeting of the nematode vesicular GABA-transporter UNC-47 (43). LAMP3 could take over a similar function, e.g., by modulating or regulating the trafficking or stability of one of the integral LB proteins like SP-C (which is synthesized as a type II transmembrane protein). However, such a mechanism might not be essential and become relevant only under conditions, where surfactant levels need to be regulated more tightly. It is interesting to note that an aggravation of the lipid changes under the pathological conditions of experimental allergic asthma may explain the lung function differences observed after MCh-provocation, when lung function was pushed to the limit. In sensitized animals, inhalation of allergen aerosols results in an acute inflammatory reaction with an infiltration of T helper 2 (TH2) cells and eosinophils into the airways and therelease of a plethora of cytokines, chemokines, and cytotoxic metabolites. Consequently, this causes activation of submucosal glands and differentiation of goblet cells, ultimately resulting in increased mucus production. TH2-type cytokines, together with eosinophil products, trigger the development of AHR that leads to an increased airway resistance in response to non-allergic stimuli and the formation of the typical asthma attack. Therefore, in mice with experimental allergic asthma, the degree of allergic airway inflammation is closely correlated with the degree of mucus production, and AHR. Remarkably, this is not the case in *Lamp3*^*-/-*^mice: While numbers of inflammatory cells such as eosinophils and lymphocytes as well as the number of goblet cells and amount of mucus stored in the airway mucosa are not different between wildtype and *Lamp3*^*-/-*^ mice, *Lamp3*^*-/-*^ mice display a markedly increased airway resistance in response to MCh inhalation. Since this response appears to be independent of both the degree of airway inflammation and mucus production, it is possible that the increase in airway resistance is not due to an increased AHR but could originate in an altered composition and thus a function of surfactant lining the alveoli and airways of *Lamp3*^*-/*-^ mice. Hence, in mice, not only the alveoli but also the *ductūs alveolares* are affected by the tension-reducing effects of surfactant. LAMP3 deficiency leads to an altered lipid composition of surfactant and consequently to a reduced effect on surface tension, which could in turn physiologically lead to increased surface tension in the distal parts of the airway system and, thus, to an increased total resistance under the fierce conditions of a MCh-provocation test.

Taken together, whereas the lack of LAMP3 seems to be compensable under physiological conditions, it impacts basal surfactant homeostasis and lung lipid composition and airway resistance in experimental allergic asthma.

## Acknowledgment

We thank Gabriele Walter, Maike Langer, Linda Lang, Juliane Artelt, Franziska Beyersdorf, and Dominika Biedziak for excellent technical assistance. We thank Dr. Nils Hoffmann (ISAS Dortmund) for support in data upload and data base management of the LIFS webportal. The authors declare no financial and non-financial competing interests. Radislav Sedlacek’s work was supported by funding of the Ministry of Education, Youth and Sports to the Czech Centre for Phenogenomics (LM2018126) and by the Institute of Molecular Genetics of the Czech Academy of Sciences (RVO 68378050). Research in Dominik Schwudke’s lab was supported by Lipidomics Informatics for Life Science (LIFS) of the German Network for Bioinformatics Infrastructure (de.NBI, BMBF).

## Author contribution

LPL, DK, DT, CS, NL, JH, MO, BS, RS, PS, DS, MW, and MD contributed to data acquisition, analysis, and interpretation. LPL, MW, and MD concepted and designed the study. MD and MW draft the initial version of the manuscript. DK and DS designed and performed all lipidomics experiments and all related data analysis. All authors were involved in drafting the final manuscript, have provided final approval of this manuscript for submission, and agreed to be accountable for their specific contribution of this study.

## Abbreviations

(AT2): Alveolar type II
(AHR): Airway hyperresponsiveness
(ANOVA): analysis of variance
(BAL): bronchoalveolar lavage
(CE): cholesterol ester
(Cer): ceramide
(CL): cardiolipin
(DAG): diacylglyceide
(DC): dendritic cells
(DPPC): dipalmitoylphosphatidylcholine
(HCA): hierarchical cluster analysis
(HexCer): hexosylceramide
(KO): knockout
(LAMP): lysosome-associated membrane glycoprotein
(LB): lamellar bodies
(LPC): lyso-phosphatidylcholine
(OVA): ovalbumin
(PC): phosphatidylcholines
(PCA): principal component analysis
(PG): phosphatidylglycerol
(LPG): lyso-phosphatidylglycerol
(PI): phosphatidylinositol
(LPI): lyso-phosphatidylinositol
(PE): phosphatidylethanolamine
(LPE): lyso-phosphatidylethanolamine
(LCL): lyso-cardiolipin
(PS): phosphatidylserine
(LPS): lyso-phosphatidylserine
(SM): sphingomyelin
(SP): surfactant protein
(TAG): triacylglyceride
(TH2): T helper cell
(TUNEL): terminal deoxynucleotidyl transferase dUTP nick end labeling

