## Supplement for "LAMP3 is critical for surfactant homeostasis in mice"

**SUPPLEMENTARY FIGURES**

**SUPPLEMENTARY METHODS**

### SUPPLEMENTARY FIGURES

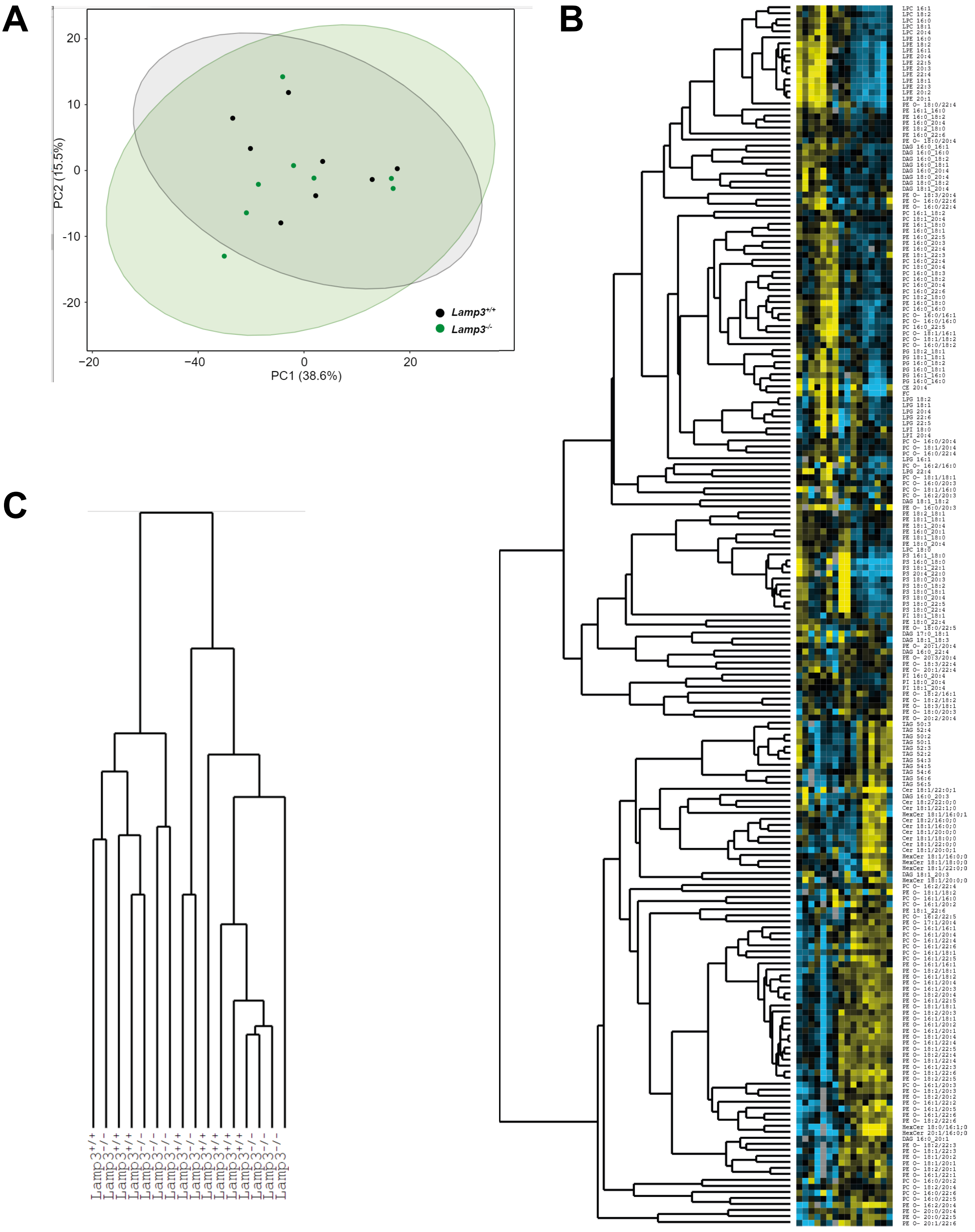


**Supplemental Figure S1. Deficiency of LAMP3 results in no observable changes in the lipidome composition under basal conditions. (A)** Two-dimensional principal component analysis (PCA) loadings plot of the BAL lipidome of wild type (black) and *Lamp3*$^{-/-}$ (green) mice. Each point represents one individual animal. Ellipses show the 95% confidence interval of the group. PC1 and PC2 explain 54.1% of the variability in the data set. Confidence intervals show a complete overlapping of samples belonging to both groups. Both groups are not separated by PCA. **(B)** Hierarchical clustering of 158 lipid species (of 393 after application of 90% occupation threshold) identified in eight wildtype and eight *Lamp3*$^{-/-}$ mice lung tissue samples. Each row represents a lipid species and each column a sample. **(C)** The hierarchical clustered tree of sample groups is not able to correlate both samples into two distinct branches.

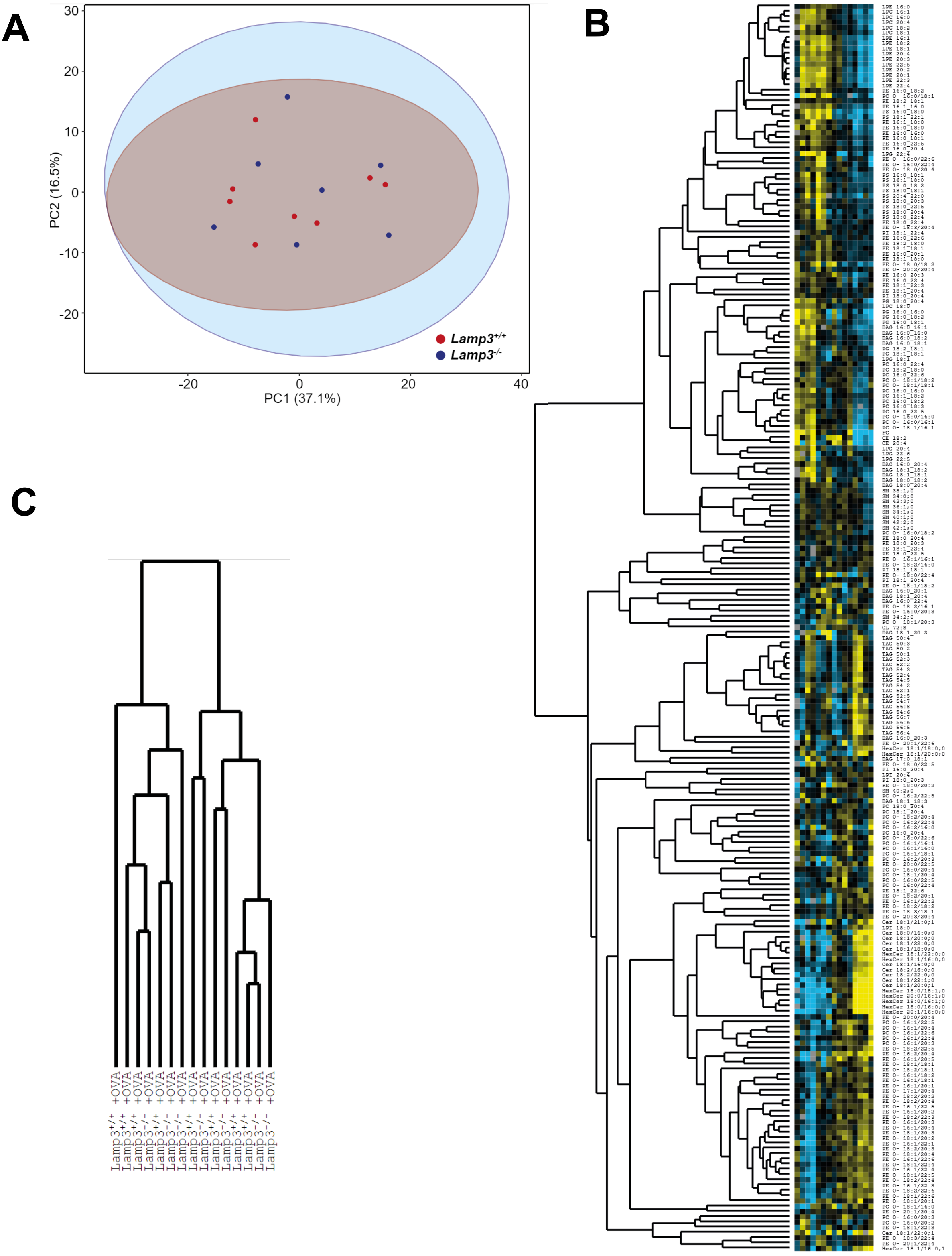


**Supplemental Figure S2. Deficiency of LAMP3 results in no observable differences in the lipidome composition when inducing the allergic asthma model. (A)** Two-dimensional principal component analysis (PCA) loadings plot of the BAL lipidome of wild type (blue) and *Lamp3*$^{-/-}$ (red) mice. Each point represents one individual animal. Ellipses show the 95% confidence interval of the group. PC1 and PC2 explain 53.6% of the variability in the data set. Confidence intervals show a complete overlapping of samples belonging to both groups. No separation of samples into two distinct groups is performed by PCA. **(B)** Hierarchical clustering of 237 lipid species (of 394 after a threshold of 90% occupation was applied) identified in eight wildtype and seven *Lamp3*$^{-/-}$ mice lung tissue samples. Each row represents a lipid species and each column a sample. **(C)** The hierarchical clustering analysis clusters samples into multiple branches not correlating to the distinct groups.

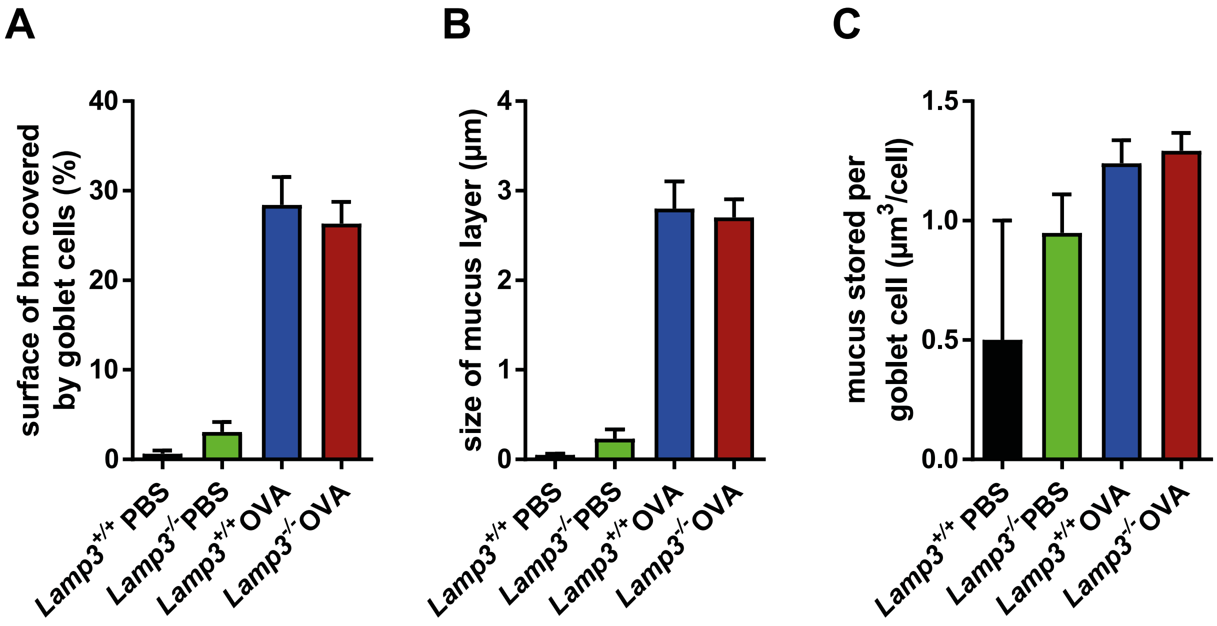


**Supplemental Figure S3. Mucus production is not altered in *Lamp3*$^{-/-}$ mice with or without OVA.** Quantification of mucus production in airway epithelial cells. OVA-induced experimental asthma increases mucus production. *Lamp3*$^{-/-}$ phenotype does not alter the amount and distribution of cells stained positive for mucus. Area of epithelial basal membrane covered by goblet cells (**A**), stored mucus volume per basal membrane area (**B**), and mucus stored per goblet cell (**C**). A-C: n = 8 mice per group.

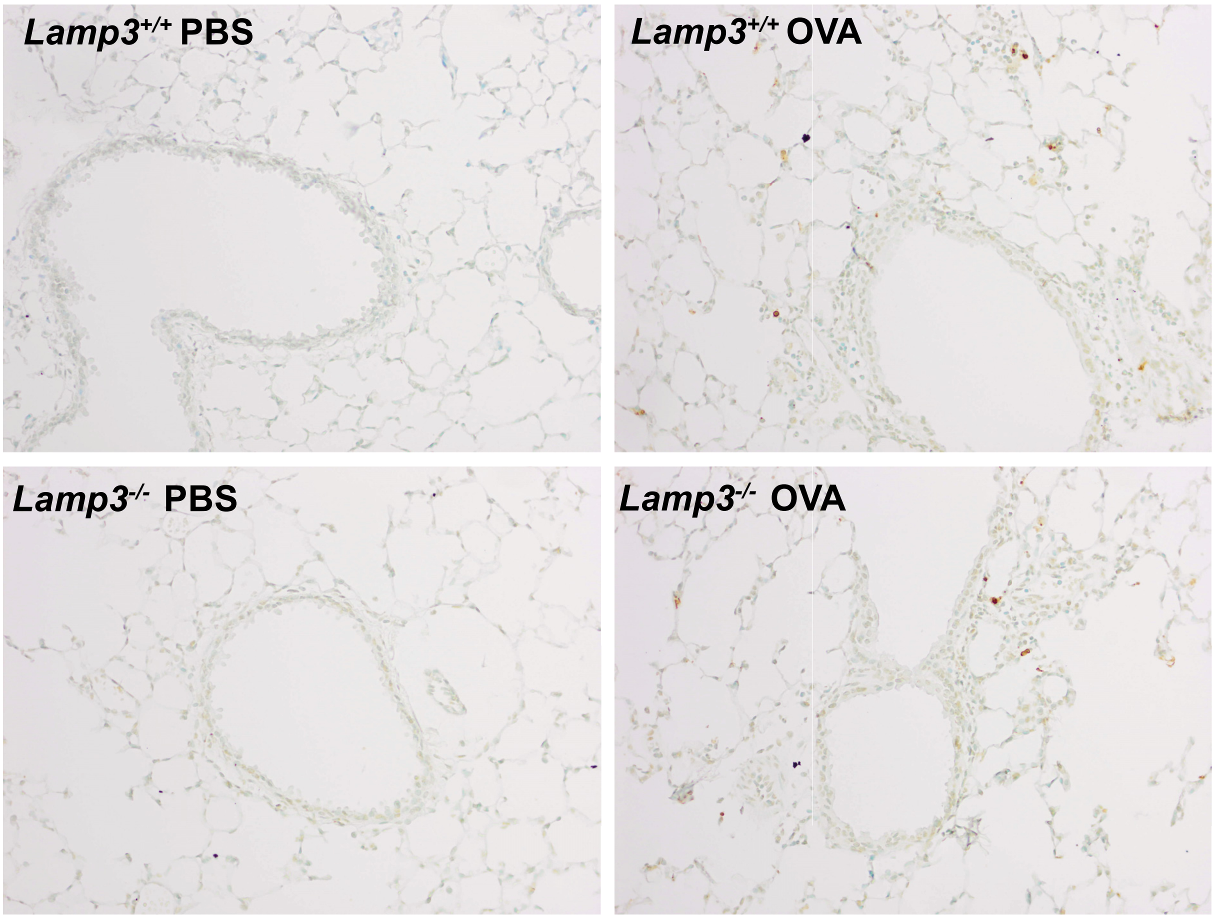


**Supplemental Figure S4. Apoptosis is not altered in *Lamp3*$^{-/-}$ mice with or without OVA.** Apoptosis staining (TUNEL assay) of inflated formalin-fixed paraffin embedded crosssections of the lung. *Lamp3*$^{-/-}$ phenotype does not alter the amount and distribution of cells stained positive for apoptosis (brown staining). Representative staining of healthy wildtype (upper left), healthy knockout *Lamp3*$^{-/-}$ (lower left), asthmatic wildtype (upper right), and asthmatic knockout *Lamp3*$^{-/-}$ (lower right).

### SUPPLEMENTARY METHODS

*Lipid extraction*

The frozen lung homogenate was diluted with 50 mM KCl solution at a constant ratio of 1:20 (wt/v), and BHT solution in methanol (1 mg/ml) was added at a ratio of 10:1 (wt/v). Samples were diluted 1:1 with 50 mM KCl solution, and 160 µl tissue homogenate containing 4 mg tissue was extracted. Further processing of 160 µl tissue homogenate and 50 µl BAL followed the same protocol.

280 µl acidified (3% v/v acetic acid) methanol and 20 µl internal standard mix A for tissue or standard mix B for BAL are added and vigorously mixed (**Table S1**). 1 ml MTBE was added, and the solution was incubated for 60 minutes at room temperature with shaking at 600 rpm. 200 µl water was added, and the samples were shaken for 20 minutes at room temperature at 1200 rpm. Afterward, the phases were separated via centrifugation at 4500 xg for 10 minutes at 15° C. 800 µl upper organic phase was transferred to a new tube, and a second extraction of the aqueous phase was performed with 500 µl saturated organic phase of MTBE/acidified methanol (+3% v/v acetic acid)/water (10:3:2.5; v/v/v)). The combined organic phases of 1300 µl were dried in a vacuum centrifuge at room temperature. The extract was resuspended in 40 µl of storage solution, aliquoted for lipidomics measurements and cholesterol derivatization, and stored at -20 °C.

*Determination of protein content*

The remaining aqueous phase of the lipid extraction was dried and afterward resuspended in 200 µl buffer (1% (wt/v) SDS buffered to pH 6,8 with Tris/HCl). Aliquots of 25 µL volume of the resulting solution were used for protein quantitation using a standard BCA kit (Thermo Fisher Scientific, Waltham, MA, USA) following the manufacturers' protocol. Absorbance was measured with an Infinite 2000 Pro plate reader (Tecan, Maennedorf, Switzerland) at 562 nm in technical triplicates.

*Cholesterol Determination*

10 μL of the BAL lipid extract and 10 μL of the tissue extract were dried in a vacuum centrifuge (Thermo Fisher Scientific, Waltham, MA, USA). Afterward, 100 μl of a solution of diluted acetylchloride in chloroform (1:5; v/v) and was added and incubated for 60 minutes at room temperature. The solution was dried in a vacuum centrifuge (Thermo Fisher Scientific, Waltham, MA, USA), resuspended in 10 μl storage solution, and subsequently stored at -20 °C until measurement.

*Internal Standards*

Standard mix A for tissue was created by combining SPLASH LIPIDOMIX Mass Spec Standard (Avanti Polar Lipids, Alabama, USA) with ceramide standard Cer d18:1/25:0 (Avanti Polar Lipids, Alabama, USA). Standard mix B for BAL was created by combining SPLASH LIPIDOMIX Mass Spec Standard (Avanti Polar Lipids, Alabama, USA) with ceramide standard Cer d18:1/17:0 (Avanti Polar Lipids, Alabama, USA). Amounts added to the samples are given in [pmol] for each internal standard aliquot are found in **Table S1**.

**Table S1: Lipid classes of the internal standard mixes are given with the final concentrations in [pmol] in the samples.**

| Lipid class | Lipid species | Standard mix A | Standard mix B |
|---|---|---|---|
| PC | PC 15:0-18:1(+[2]H7) | 106.7 | 106.7 |
| PE | PE 15:0-18:1(+[2]H7) | 4.0 | 4.0 |
| PS | PS 15:0-18:1(+[2]H7) | 2.7 | 2.7 |
| PG | PG 15:0-18:1(+[2]H7) | 19.0 | 19.0 |
| PI | PI 15:0-18:1(+[2]H7) | 5.4 | 5.4 |
| PA | PA 15:0-18:1(+[2]H7) | 5.4 | 5.4 |
| LPC | LPC 18:1(+[2]H7) | 24.1 | 24.1 |
| LPE | LPE 18:1(+[2]H7) | 5.5 | 5.5 |
| CE | CE 18:1(+[2]H7) | 270.5 | 270.5 |
| MAG | MAG 18:1(+[2]H7) | 2.8 | 2.8 |
| DAG | DAG 15:0-18:1(+[2]H7) | 8.0 | 8.0 |
| TAG | TAG 15:0-18:1(+[2]H7)-15:0 | 35.3 | 35.3 |
| SM | SM d18:1-18:1(+[2]H9) | 20.9 | 20.9 |
| Chol | Cholesterol(+[2]H7) | 125.0 | 125.0 |
| Cer | Cer d18:1/17:0 | 4.71 | - |
| Cer | Cer d18:1/25:0 | - | 24.38 |

*Lipidomics data acquisition:*

Lipidomics acquisitions consisted of a total duration of 10 minutes with a switch from positive ESI mode to negative ESI mode after 5 minutes. In positive and negative mode, the first 30 seconds were used for electrospray stabilization. Afterward full MS from 350 to 1100 m/z was acquired with a following full MS scan every 100 MS² scans for a total of eight full MS scans for each ESI mode in the span of 5 minutes. Full MS acquisitions were performed with a micro scan count of five, a max injection time of 200 ms, and a targeted resolving power of 2.8e5. MS² acquisitions were performed on parent ions from 350 to 1200 m/z with an inclusion list. The lower mass in MS² acquisitions was kept at 180 in positive mode and 150 in negative mode with the upper mass scaling to the parent ion. MS² acquisitions were performed with a micro scan count of one, a targeted resolving power of 7e4, a MS² isolation width of 1 m/z, a max injection time of 200 ms, an AGC target of 1e5, and a HCD energy of 30 eV.

*Free cholesterol data acquisition:*

Cholesterol acquisitions consisted of a total duration of two minutes in positive ESI mode. MS was acquired for the mass window of 440 to 460 m/z followed by $MS^2$ acquisitions of 446.3993@hcd15.00 [250-475] and 453.4432@hcd15.00 [250-480]. MS acquisitions were performed with a micro scan count of one, a targeted resolving power of 2.8e5, a max injection time of 200 ms, an AGC target of 3e6. $MS^2$ acquisitions are performed with a micro scan count of one, a targeted resolving power of 2.8e5, a max injection time of 100 ms, an AGC target of 2e5 a $MS^2$ isolation width of 1 m/z, and a HCD energy of 15 eV.

*MS Data extraction:*

The following options were supplied to msconvert for shotgun lipidomics acquisitions:

[--32 --zlib --filter "peakPicking vendor msLevel=1-" —filter "titleMaker <RunId>.<ScanNumber>.<ScanNumber>.<ChargeState> File:<SourcePath>, NativeID:<Id>]

Options for derivatized cholesterol measurements are as follows:

[--32 --zlib --filter "peakPicking vendor msLevel=1-" --filter "zeroSamples removeExtra" –simAsSpectra]

*Lipid identification with LipidXplorer*

MS information was selected with a selection window of 0.4 Da, a resolution of 2.4e5 at m/z 350, a tolerance of 4 ppm, a relative threshold of 0.001%, a resolution gradient of -125 with a minimum occupation of 0.01 and without a frequency filter and MS offset. $MS^2$ information was selected with a resolution of 9e4, a tolerance of 4 ppm, a relative threshold of 0.001%, a resolution gradient of -80, minimum occupation of 0.1, and without a frequency filter or a PMO. Lipid species identifications were assigned when the mass accuracy was ≤3 ppm. The lipid classes PC, PC O-, LPC, PE, PE O-, LPE, PA, PI, LPI, PG, LPG, PS, LPS, CL, LCL were identified in negative ESI mode and TAG, DAG, SM, CE, Cer and HexCer in positive ESI mode via customized MFQL scripts. Derivatized cholesterol is identified with a tolerance of 5 ppm. Custom MFQL scripts, data and settings can be downloaded from https://apps.lifs.isas.de/.

*Post-processing and Quantification of lipids with lxPostman*

Output files from LipidXplorer were loaded into lxPostman and merged. False-positive signals introduced by the extraction process were removed when lipid species were less than 10 times higher than in control blank samples. Lipid species-specific fragments were specified in the MFQL files, and fragment intensities were used to determine intensity ratios. Fragmentation ratios were computed for the acyl fragment ions (fragment A / B) for PA, PC, PE, CL, PG, PI, PS and for precursor and acyl fragment (parent / fragment A) for LPC and LPE in the negative ion mode. For DAG quantified in the positive ion mode, the intensity ratio of the neutral loss of the fatty acids (fragment A / B) was utilized. The fragmentation ratios of internal standards were used to filter eligible ratios within a range of 3 and 1/3 times the standard value. The quantification is based on the intensity ratio to internal standards (Table S1) of known concentration. A minimum occupation threshold for each lipid species across all samples is set to 5% to filter sporadic occurring lipid species but keep lipid species present in multiple samples. A regularly actualized version of lxPostman can be found at https://apps.lifs.isas.de/ in the tab "R shiny Apps".
